## Supplementary figures and images for "Dynamic Turnover of PI4P and PI(4,5)P2 under Hypoxia Controls Electrostatic Plasma Membrane Targeting"

### Figure_S1.jpg

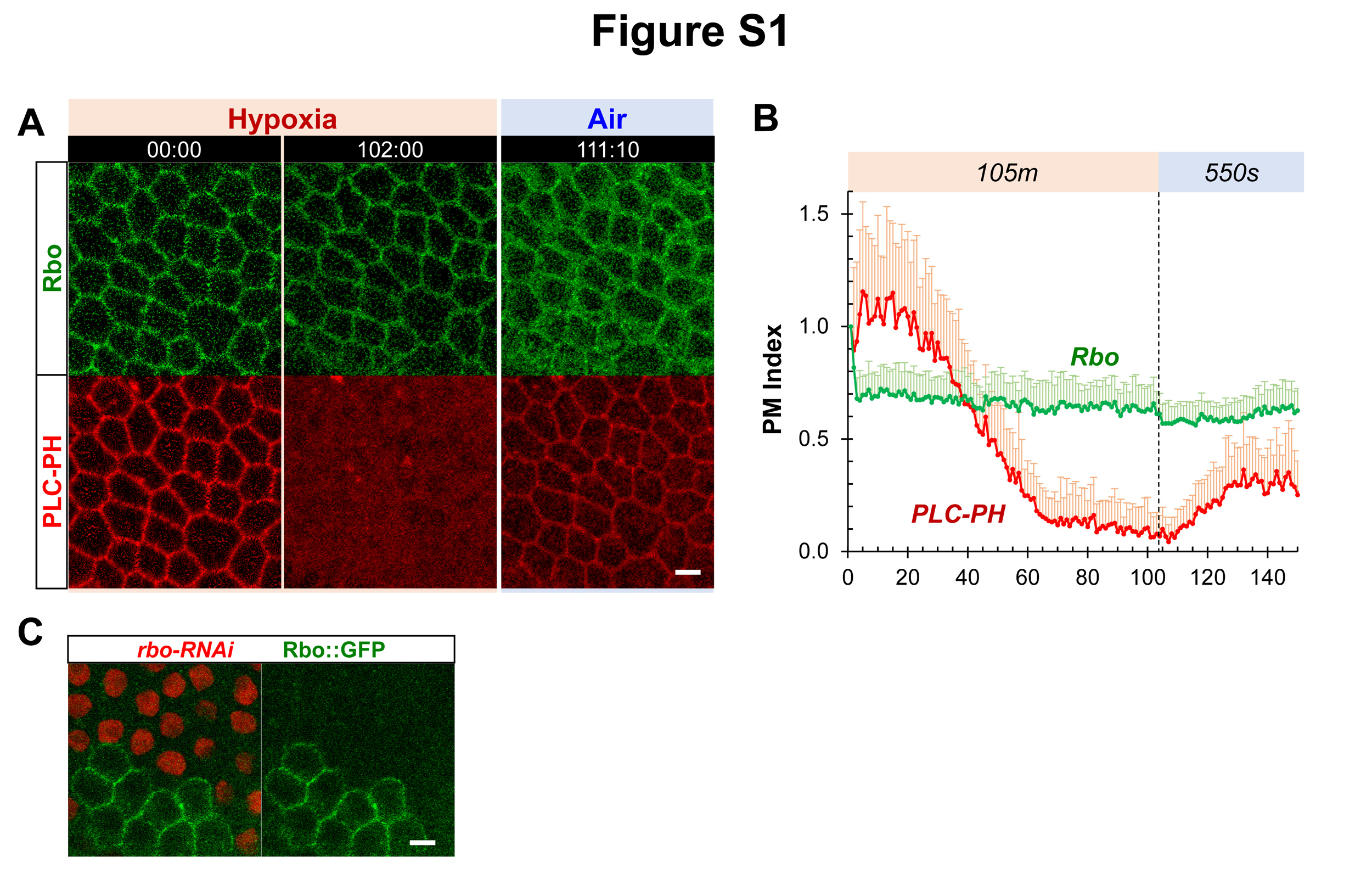

### Figure_S2.jpg

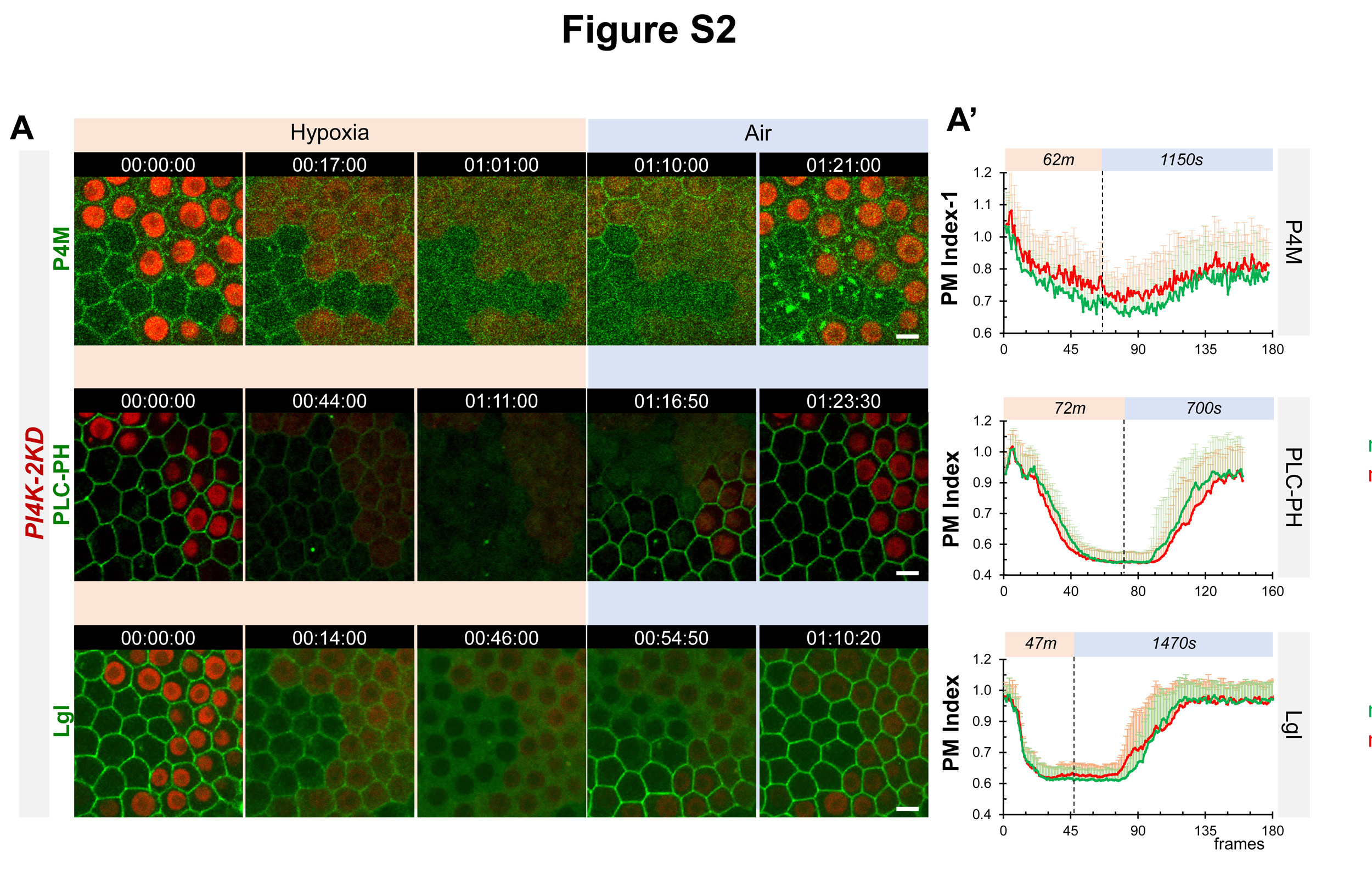

### Figure_S3.jpg

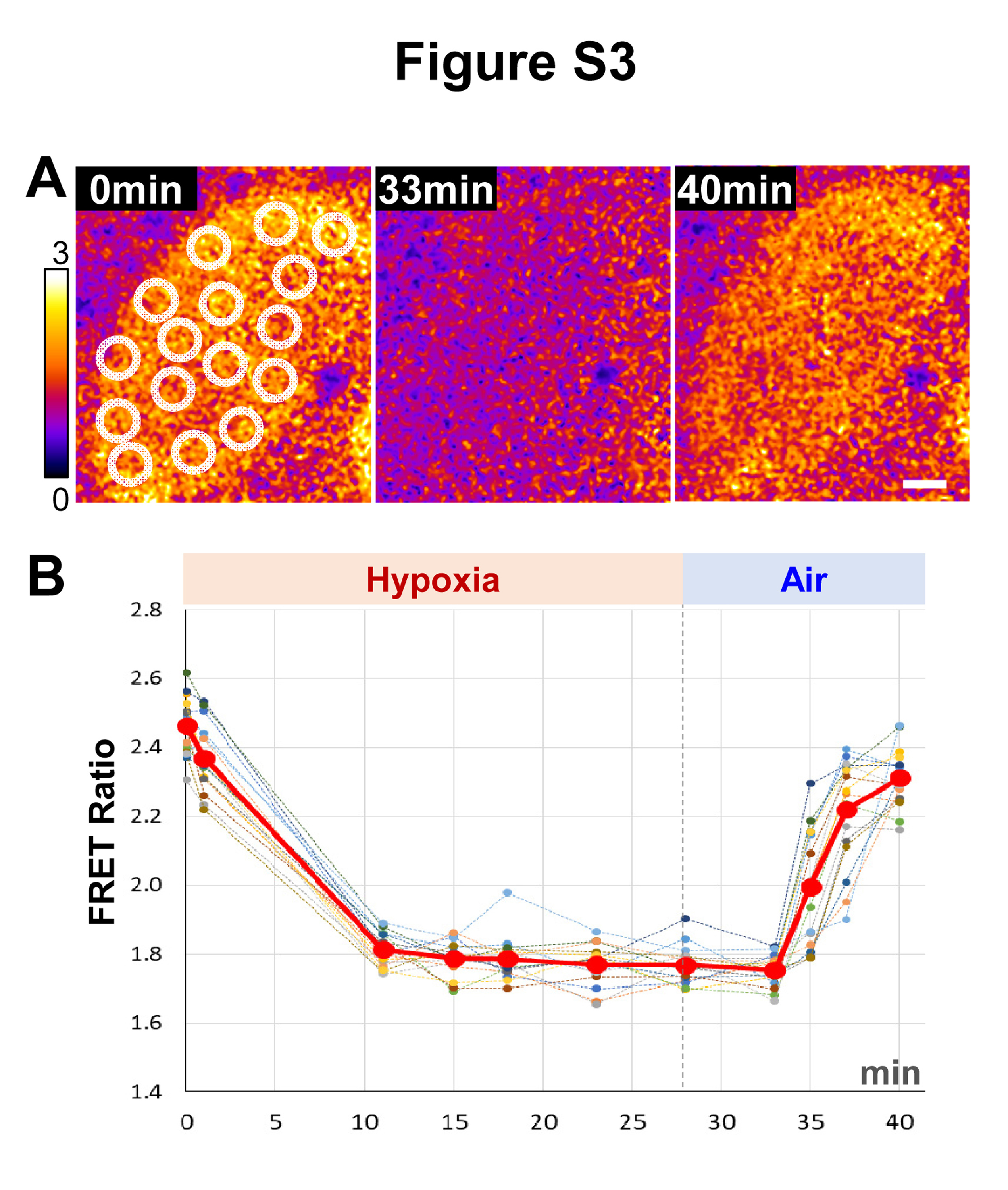

### Figure_S4.jpg

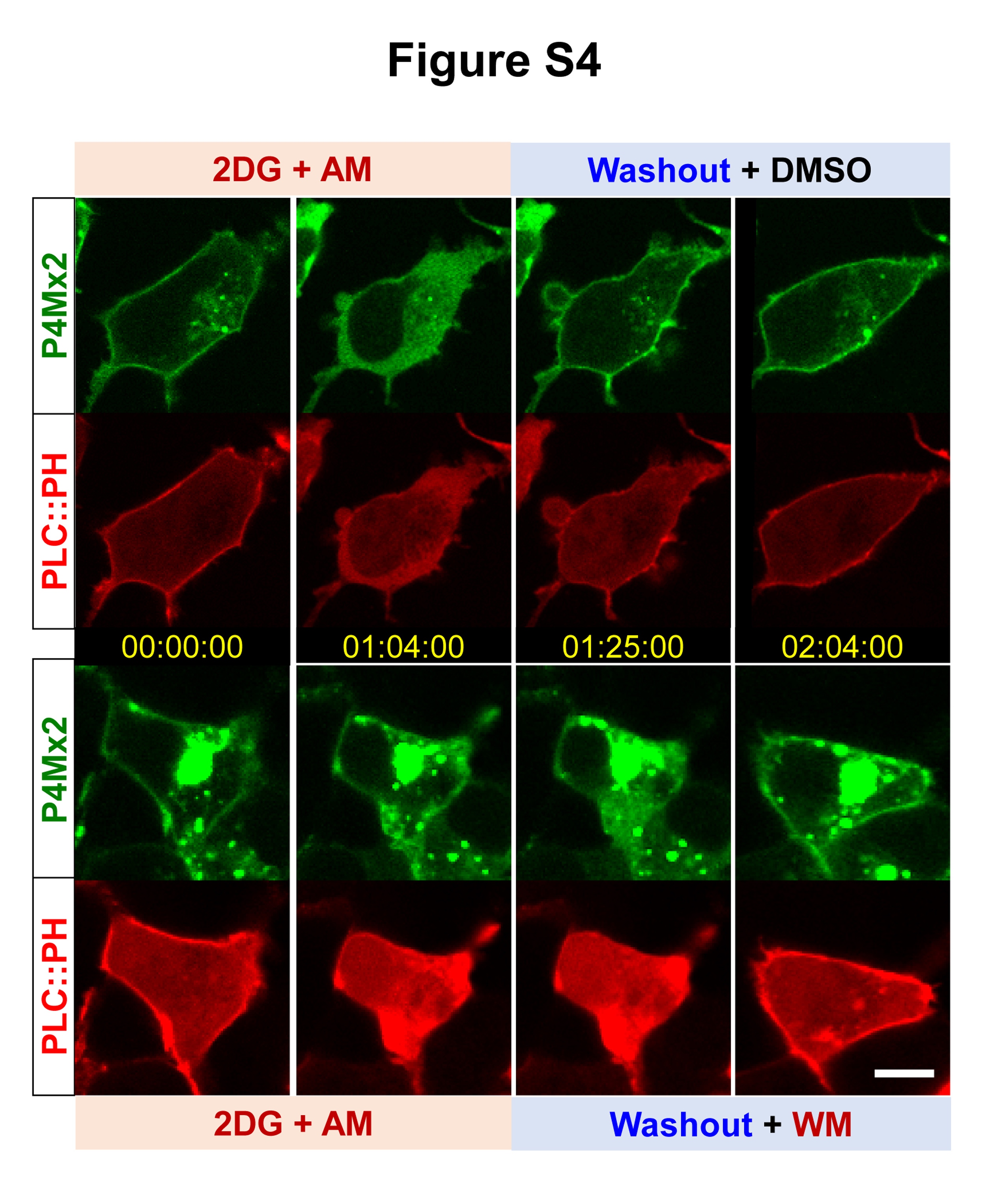
